# A new causal centrality measure reveals the prominent role of subcortical structures in the causal architecture of the extended default mode network

**DOI:** 10.1101/2023.04.22.537911

**Authors:** Tahereh S. Zarghami

## Abstract

Network representation has been a groundbreaking concept for understanding the behavior of complex systems in social sciences, biology, neuroscience, and beyond. Network science is mathematically founded on graph theory, where nodal importance is gauged using measures of *centrality*. Notably, recent work suggests that the topological centrality of a node should not be over-interpreted as its dynamical or causal importance in the network. Hence, identifying the influential nodes in dynamic causal models (DCM) remains an open question. This paper introduces *causal centrality* for DCM, a dynamics-sensitive and causally-founded centrality measure based on the notion of *intervention* in graphical models. Operationally, this measure simplifies to an identifiable expression using Bayesian model reduction. As a proof of concept, the average DCM of the extended default mode network (eDMN) was computed in 74 healthy subjects. Next, causal centralities of different regions were computed for this causal graph, and compared against major graph-theoretical centralities. The results showed that the *subcortical* structures of the eDMN are more causally central than the *cortical* regions, even though the (dynamics-free) graph-theoretical centralities unanimously favor the latter. Importantly, model comparison revealed that only the pattern of causal centrality was *causally relevant*. These results are consistent with the crucial role of the subcortical structures in the neuromodulatory systems of the brain, and highlight their contribution to the organization of large-scale networks. Potential applications of causal centrality - to study other neurotypical and pathological functional networks – are discussed, and some future lines of research are outlined.

## 1 Introduction

Network analysis has found widespread application in different disciplines including sociology, psychology and neuroscience (Barabási 2014; Bassett and Sporns 2017; Borsboom and Cramer 2013; Marsman et al. 2018; McNally 2016; Otte and Rousseau 2002). One of the foundational tools in network science is graph theory, a branch of mathematics that examines the properties of networks represented as graphs^1^. A crucial application of graph theory is to gauge the relative importance (aka *centrality*) of individual nodes in a network. One promising prospect is for the centrality information to guide clinical interventions in the network and to predict the outcomes thereof (Gessell et al. 2021).

Numerous centrality measures have been developed based on different definitions for nodal *importance*; degree, closeness and betweenness centralities are prominent examples (Freeman 1979; Opsahl et al. 2010). Notably, centrality measures are based only on the network *topology*, not *dynamics*—even when a flow process is implicitly associated (Klemm et al. 2012). Importantly, if the flow assumptions of a centrality measure do not match the flow characteristics of the network to which it is applied, the centrality ranking of the nodes can be simply “wrong” (Borgatti 2005). Moreover, recent work has shown that graph-theoretical (GT) centrality measures are “poor substitutes” for the *causal influence* of the nodes (Dablander and Hinne 2019). Instead, the latter should be estimated using dedicated methods in the field of *causal inference* (Glymour et al. 2016).

Causal inference is a growing discipline that aims to answer *causal* rather than associative questions. A well-known framework for representing causal or directional relationships is directed graphical models, which consists of directed *acyclic* graphs (DAG) and directed *cyclic* graphs (DCG) (Glymour et al. 2016; Pearl 1998). Compared to DAGs, DCGs have been much less characterized (Ghassami et al. 2020; Park and Raskutti 2016; Pearl 1998; Richardson 1996a, 1996b; Spirtes 1995), even though cyclic effects are inevitable in many applications. Specifically, in the brain, the reciprocal polysynaptic influences of neuronal populations create feedback (cyclic) loops. Hence, a class of time-dependent DCGs has been particularly developed to represent dynamical models of causal influences in the brain— known as dynamic causal models (DCM). These are state-space models that govern the dynamics of coupled neuronal states and their mappings to observed signals (such as neuroimaging data) (Friston et al. 2003).

The distinction between structural and dynamic causal models has been schematically illustrated in Fig. 1, based on the detailed comparison in (Friston 2011). Briefly, structural causal models (SCM) are based on DAGs and static nonlinearities, whereas DCM accommodates time-dependent DCGs. In the DCM graph, nodes represent hidden neuronal states and directed edges encode the reciprocal neuronal couplings (aka *effective connectivity* parameters). The hidden states are mapped to observed variables through an observation function (Fig. 2). Given empirical observations, the generative model of DCM is inverted using variational Bayesian inference to estimate the optimal model parameters and the associated model evidence (Friston et al. 2007). Once the causal effects have been estimated, a natural question arises: what is the relative significance of each node (i.e., brain region) in this DCM?

**Fig. 1:**
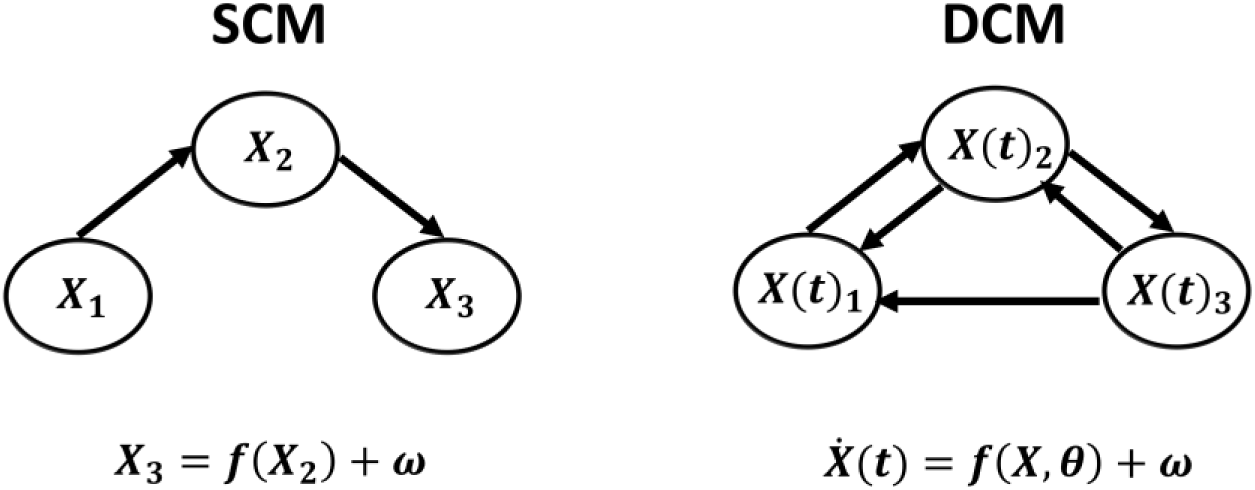
Structural and dynamic causal modeling. (Left): SCM is mainly concerned with discovering conditional independencies induced by static nonlinear mappings, in DAGs. (Right): DCM is based on time-dependent DCGs, expressed as differential equations. For simplicity, the mappings from hidden neuronal states to the observations are not shown for this DCM (see Fig. 2 instead). Abbreviations: DAG, directed acyclic graph; DCG, directed cyclic graph; SCM, structural causal model; DCM, dynamic causal model or directed cyclic model.

**Fig. 2:**
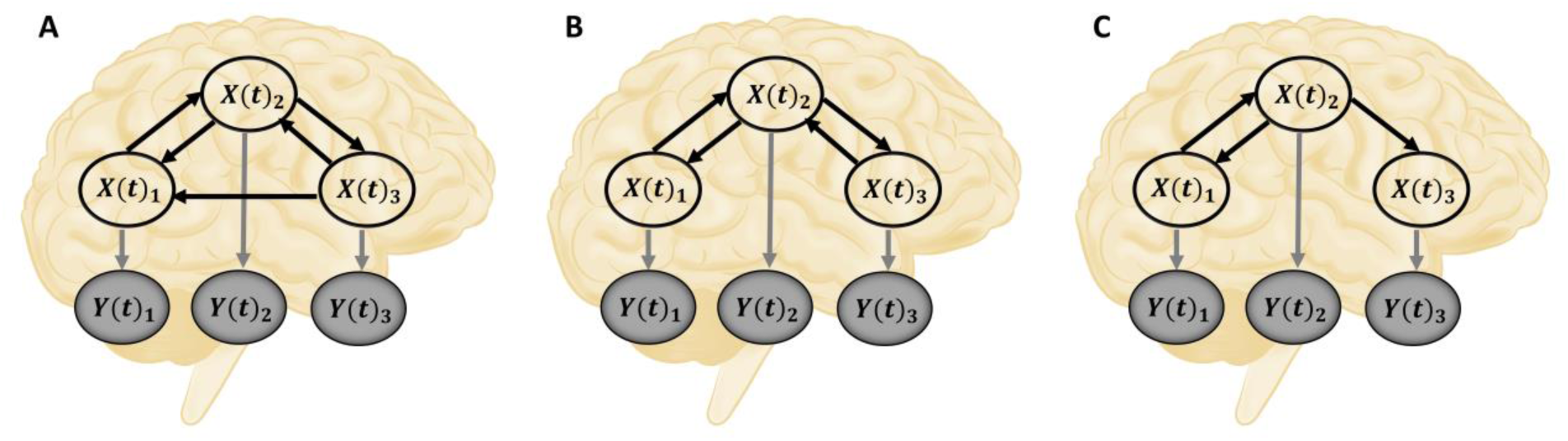
Intervention on the edges of a DCM. (A) Pre-intervention graph of an exemplar DCM with three hidden states. *X*-nodes represent the hidden neuronal states and *Y*-nodes (shaded) denote the observed variables. Edges represent the causal influences. (B) Post-intervention graph, after intervention on (i.e., removing) the *X*_3_ → *X*_1_ edge. (C) Post-intervention graph, after intervention on all the causal influences of *X*_3_ directed at the other hidden states. (Brain schematic: Injurymap, <u>Human Brain</u>, Recolor by Author, <u>CC BY 4.0</u>).

Recently, (Dablander and Hinne 2019) showed that the graph-theoretical centrality of a node is *not* indicative of its causal importance in a DAG. To quantify the latter, the authors defined the *total causal effect* of a node as the sum of the node’s causal influences over its children in the DAG (Dablander and Hinne 2019). However, in a cyclic graph like a DCM, a node’s children might as well be its parents. So, how can one quantify the causal and dynamical significance of the nodes in a DCM?

To answer to this question, a *causal centrality* measure for DCM is introduced in this paper, based on the notion of *intervention* in graphical models. We will see that this measure is *identifiable* by virtue of recent developments in Bayesian (sub)model comparison—known as *Bayesian model reduction* (BMR). Then, through a worked example, we shall compare causal centrality with some common graph-theoretical centralities, and asses the *causal relevance* of each measure in a model comparison framework. In the following, some background is reviewed before the new measure is introduced.

## 2 Theory

### 2.1 Intervention in causal models

Pearl’s theory of causation, called structural causal modeling (SCM), combines features of structural equation models (SEM, used in economics and social sciences) and graphical models (developed for probabilistic and causal reasoning) (Glymour et al. 2016; Pearl 1998). While graphical models encode the causal assumptions in SCM (Fig. 1-A), causal derivations are performed in a dedicated algebraic language called the *calculus of interventions* or *do-calculus*.

*Intervention* on a node (*X*) in a graphical model, denoted as *do*(*X* = *x*_0_), deletes the edges pointing to that node, and assigns a constant value to the node *X* = *x*_0_. In this calculus, the statement *P*(*Y*|*do*(*X* = *x*_0_)) describes how the distribution of *Y* changes under the *hypothetical* setting of *X* to *x*_0_. This *post-intervention* distribution is different from the familiar conditional probably statement *P*(*Y*|*X* = *x*_0_), which describes how the distribution of *Y* changes when *X* is *observed* to be equal to *x*_0_. Using do-calculus, Pearl defined a measure of causal influence, called *average causal effect* (ACE) (Glymour et al. 2016; Pearl 2010):

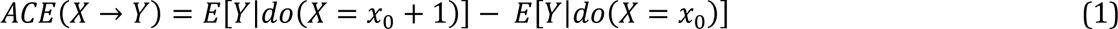

where *E* is the expectation operator with respect to the post-intervention distributions. He also showed that, under certain conditions, hypothetical quantities such as *P*(*Y*|*do*(*X* = *x*_0_)) are identifiable (i.e., estimable) from observed data (i.e., pre-intervention distributions). Total ACE of a node (*X*_*j*_) in a DAG can be computed by adding up the node’s absolute ACE values over its children (*Ch*(*X*_*j*_)) (Dablander and Hinne 2019):

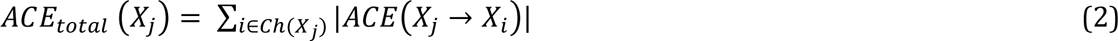

Note that ACE only accounts for *linear* dependencies between cause and effect, and it is not designed to detect whether *X* changes higher order moments of the distribution of *Y* (Eq. 1). For this and other reasons, (Janzing et al. 2013) defined an alternative measure to quantify causal strength between two nodes, using intervention on the *edges* and by measuring the Kullback-Leibler (KL) divergence between the pre- and post-intervention distributions of a DAG over its *n* nodes:

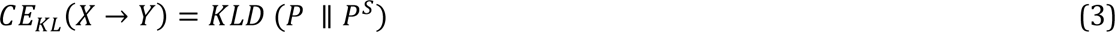

The post-intervention distribution (*P*^*s*^) corresponds to a DAG in which the directed edge (i.e., causal dependence) between *X* and *Y* is removed, and instead the marginal distribution of *X* is fed into *Y* (Janzing et al. 2013). Similarly, the *total* KL-based causal effect of a node (*X*_*j*_) in a DAG can be computed as the KL-divergence between the DAG’s pre- and post-intervention distributions, where the intervention is simultaneously applied to all the outgoing edges from that node (Dablander and Hinne 2019).

Note that both ACE and CE_KL_ quantify *causal strengths* in a DAG. The present paper tackles a related but distinct problem, which is quantifying the overall causal importance of individual nodes in a DCM, *after* the causal strengths have been estimated^2^. Still, the concepts of *intervention on the edges* and *KL-based distance* are fundamental to the proposed centrality measure. Before introducing this measure, it is worthwhile to review a Pearlian guideline for new developments in the causal framework. When discussing the possibility of extending SEMs beyond modeling linear causal dependencies, Pearl asserts that a key requirement for any such extension is to “redefine ‘effect’ as a general *capacity to transmit changes among variables* [emphasis added]. Such an extension, based on simulating hypothetical *interventions* [emphasis added] in the model […] has led to new ways of defining and estimating causal effects” (Pearl 2010). As we shall see, causal centrality is based on these guidelines.

### 2.2 Definition of causal centrality

The *causal centrality* of a node in a DCM is defined as the (normalized) KL-divergence between the pre- and post-intervention distributions of the DCM; where the distribution is the joint probability distribution over the model’s unobservable^3^ (θ) and observable (*Y*) variables, given the generative model of DCM (*m*); and intervention refers to removing the causal influences of that node directed at the other hidden nodes (Fig. 2). Mathematically:

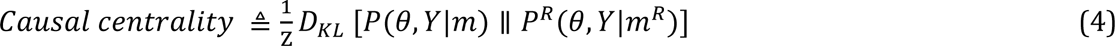

where *P* and *P*^*R*^ refer to the pre- and post-intervention distributions, associated with the *full* and *reduced* models (*m* and *m*^*R*^); and the normalization constant *Z* is the model evidence (as shown in Appendix A). In the reduced model (*m*^*R*^), some of the edges (θ_*i*_) of the full model have been removed; i.e., forcefully set to zero, in the spirit of: *P*(θ, *Y*|*do*(θ_*i*_) = 0, *m*). An example of such intervention on the edges of a DCM has been illustrated in Fig. 2. The key issue here is identifiability of the post-intervention distribution. “The central question in the analysis of causal effects is the question of identification: Can the controlled (post-intervention) distribution *P*(*Y* = *y*| *do*(*x*)) be estimated from data governed by the pre-intervention distribution, *P*(*z*, *x*, *y*)?” (Pearl 2010). As we will shortly see, Bayesian model reduction can facilitate the computation of post-intervention distributions from pre-intervention quantities.

### 2.3 Bayesian model reduction

BMR refers to the analytic inversion of a reduced model based on the priors and posteriors of the full model and the priors of the reduced model. Reduced models are nested within the full model; that is, they include only a subset of the parameters of the parent model by “switching off” the other parameters. This is achieved by imposing very precise null priors over the selected parameters, which shrinks them to zero (Friston et al. 2016; Friston et al. 2018; Friston and Penny 2011). To see how BMR can be applied in the current context, note that Eq. 4 can be rearranged as follows (details in Appendix A):

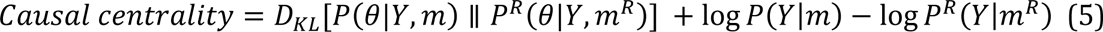

BMR facilitates the computation of model evidence (*P*^*R*^(*Y*|*m*^*R*^)) and parameter posteriors (*P*^*R*^(θ|*Y*, *m*^*R*^)) for the reduced model, using the same quantities already estimated for the full model. Specifically, when the inference is variational, we can write (see Appendix B):

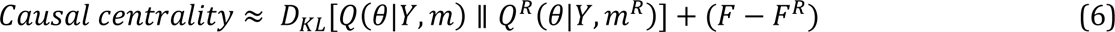

The approximate posterior (*Q*) and the free energy^4^ (*F*) of the full model are already available by virtue of the original model inversion. Hence, BMR can analytically compute *Q*^*R*^and *F*^*R*^ for the reduced (intervened upon) model (Appendix B). Thereafter, causal centrality is computed as the KL-divergence between the approximate posteriors of the full and reduced models (*Q*, *Q*^*R*^), plus the approximate change (decrease^5^) in free energy following the intervention. The Gaussian distributions assumed in variational Laplace (Friston et al. 2007; Zeidman et al. 2022) further simplify the computation of the KL-divergence (Appendix A). Note that, Eq. 6 also reveals the intuitive properties of a node with high causal centrality; that is, intervention on the causal effects of a highly central node would result in substantial drop in model evidence (*F* − *F*^*R*^), and the remaining causal strengths should be substantially revised to explain the same data *Y* (*D*_*KL*_ term).

So far, we defined causal centrality; derived an identifiable expression for its computation; and mentioned the intuition behind it. In the following, some face and construct validation is provided for this measure through a worked example. Namely, causal centralities are computed for the nodes in a DCM group study, which are next compared against several graph-theoretical centrality measures. Lastly, all these measures shall be included in a model comparison framework, to reveal the extent of their causal relevance.

## 3 Materials and Methods

This section includes an illustrative example to showcase the utility of causal centrality. This example is based on recent characterizations of the default mode network (DMN) as a functional network that extends beyond the cortical regions, into the subcortex. First, the average causal architecture of this extended DMN is identified in a group of healthy subjects; then, the graph-theoretical and causal centralities of the constituent regions are computed and contrasted against each other. Finally, each of these centrality measures is incorporated as prior information in a model comparison framework, which shall elucidate their causal relevance.

### 3.1 Dataset and pre-processing

Resting state fMRI scans from 74 healthy individuals (age: 35.8 ± 11.6; 51 males) were acquired from the publicly available COBRE dataset^6^ (Çetin et al. 2014). The subjects had been scanned for five minutes on a 3-Tesla Siemens scanner, while fixating on a central cross. A total of 150 functional volumes had been collected using gradient-echo EPI sequence, with the following settings: TR = 2 s, TE = 29 ms, flip angle = 75°, 33 axial slices, ascending acquisition, matrix size = 64 × 64, voxel size = 3.75 × 3.75 × 4.55 mm, field of view = 240 mm. High-resolution T1-weighted structural images had also been acquired for all subjects.

The data were preprocessed in SPM12^7^ as follows: the first 5 volumes were discarded to allow for T1 equilibration; the remaining functional images were realigned to the first volume, slice-timing corrected, co-registered to the structural image, normalized to the standard MNI152 template (Collins et al. 1998), resampled to 3 *mm*^3^ isotropic voxels, and smoothed with a Gaussian kernel (FWHM = 6 mm). For each subject, the absolute head motion was below one voxel, and the mean framewise displacement (MFD) was 0.35 ± 0.18 mm across the group (Power et al. 2012).

### 3.2 Extended default mode network

The default mode network (Buckner and DiNicola 2019; Raichle et al. 2001) is a collection of brain region that are typically deactivated during external goal-directed tasks, but activated during internally focused cognitive processes such as mind wandering, memory retrieval, self-referential and emotional processing, perspective taking, etc. (Knyazev et al. 2020; Murphy et al. 2018; Mwilambwe-Tshilobo and Spreng 2021; Qin and Northoff 2011). Most of these psychological functions are readily carried out during wakeful *resting state*, the context in which DMN was first characterized^8^ (Greicius et al. 2003; Raichle et al. 2001).

To date, DMN has largely been a cortically defined network, consisting of regions distributed across the ventromedial and lateral prefrontal cortex, posteromedial and inferior parietal cortex, as well as the lateral and medial temporal cortex. As such, DMN has been considered a backbone of cortical integration (Alves et al. 2019; Andrews-Hanna et al. 2010; Kernbach et al. 2018; Lopez-Persem et al. 2019; Margulies et al. 2016). Hence, the involvement of subcortical regions in the DMN has been usually overlooked. Importantly, subcortical structures contain the neurochemically diverse nuclei that are crucial in the pathophysiology of various brain diseases, in which DMN connectivity is gravely affected; these include Alzheimer’s disease, Parkinson’s disease, schizophrenia, depression and temporal lobe epilepsy, among others (Kottaram et al. 2019; Mothersill et al. 2017; Parsons et al. 2020; Qian et al. 2019; Ruppert et al. 2021; Sheline et al. 2009; Whitfield-Gabrieli and Ford 2012). Therefore, investigating the contribution of subcortical structures to the DMN function is an essential step towards understanding the mechanisms of DMN disruption in these disorders, and more broadly serves to reconcile neurochemistry, connectivity and cognition.

Recently, (Alves et al. 2019) have revisited the spatial extent of DMN, with specific focus on the structural and functional role of the subcortical regions in the DMN integration. The result has been an improved neuroanatomical model of the DMN, consisting of 33 cortical and subcortical regions of interest (ROIs). These regions and their MNI coordinates have been outlined in Table 2 of (Alves et al. 2019), and the final template has been shared on the NeuroVault^9^ repository. This extended DMN (eDMN) was used in the present work as prior spatial information to identify subject-specific eDMNs in 74 healthy subjects, which were then causally modeled using DCM—as elaborated shortly.

### 3.3 Identifying eDMN

To identify the extended DMN for each subject, the eDMN template was used as prior information in a spatially-constrained independent component analysis (SC-ICA) (Lin et al. 2010). Constrained ICA is a semi-blind source separation method, where prior information is added to the contrast function of a standard blind ICA in the form of (in)equality constraints. As such, the prior information ensures some level of similarity between the recovered sources and the reference templates, and keeps the optimization solution bounded. Importantly, SC-ICA accommodates the spatial variability of the networks between subjects (unlike fixed templates) while maintaining the component (i.e., network) correspondence over the group. SC-ICA algorithm has been implemented in the Group ICA of fMRI Toolbox (GIFT^10^).

To assess the consistency and fidelity of the identified networks, the validation procedure in (Lin et al. 2010) was followed. Briefly, voxel-wise one sample t-tests were performed across all the eDMNs (after variance normalizing each spatial pattern). The resulting t-map was thresholded at a false discovery rate (FDR) corrected q-value < 0.01, to create a group eDMN map. Subject-specific eDMNs were then intersected with this group eDMN to yield group-adjusted individual networks. Finally, the normalized spatial correlation of the template with each adjusted individual eDMN was computed, which turned out to be 0.81 ± 0.03 across the group. In short, this procedure revealed the fidelity of individual networks to the prior template, as well as natural spatial variability over the subjects. The template from (Alves et al. 2019) and the average spatial map of eDMN (from the present study) have been illustrated in Fig. S1 and Fig. S2.

### 3.4 Dynamic causal modeling the eDMN

#### 3.4.1 Region specification and time series extraction

Several eDMN regions were merged based on physical and functional proximity for computational tractability of the DCM analysis^11^. First, the 16 pairs of highly correlated homologous regions were merged. Next, VMPFC^12^, AMPFC^13^ and DPFC^14^ were grouped into one prefrontal (PFC) component; two temporal sub-regions (TP^15^ and MTG^16^) were merged into one temporal (Tmp) component; and two cerebellar regions (CbH^17^ and CbT^18^) were represented by one cerebellar (Cb) component. This resulted in 13 regions for spectral dynamic causal modeling, which have been listed in Table 1.

**Table 1:**
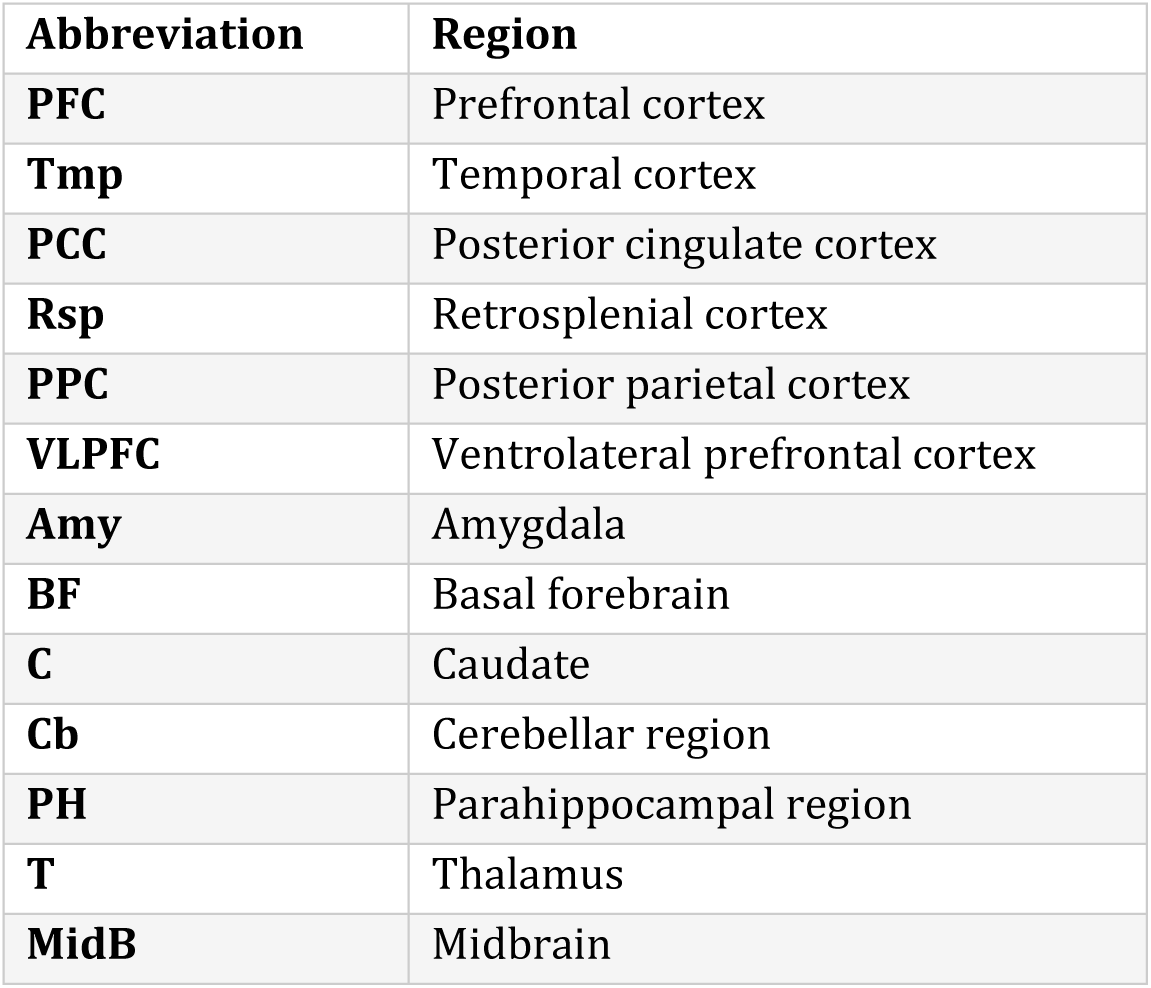
The regions of eDMN included in DCM analysis.

The average regional time series were extracted for these 13 ROIs from the intersection of group-adjusted individual eDMNs with the parcellation map of (Alves et al. 2019). Post-processing (Allen et al. 2014; Damaraju et al. 2014; Iraji et al. 2022) of the time series included: removal of the linear trends, regression of the nuisance (motion and physiological) signals, and replacement of outlier time points with spline interpolations (using AFNI’s 3dDespike). The average functional connectivity between these regions has been depicted in Fig. S3. The post-processed time series were next modeled using spectral DCM.

#### 3.4.2 Spectral dynamic causal model

Spectral DCM was used to estimate the effective (causal) connections among the 13 regions of eDMN, for each subject. The generative model of this DCM specifies how the observed complex cross-spectra^19^ of fMRI signals are generated from the interaction of coupled neuronal ensembles and the neurovascular coupling. The model is initially expressed in the time domain as a state-space model with two equations; the first equation describes the dynamics of an endogenously driven network of coupled neuronal populations, and the second equation maps these neuronal activities to the observed hemodynamic responses (Friston et al. 2014; Razi et al. 2015):

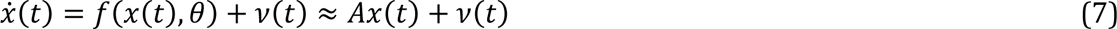

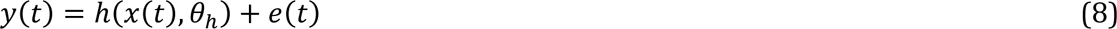

where *x*(*t*) is a vector containing the hidden neural states (i.e., ensemble activities) of *n* brain regions, at time *t*. The rates of change^20^ in neural states *x*(*t*) are influenced by the activities of (self and) other regions via the connectivity matrix, *A*, and fast endogenous neural fluctuations, *v*(*t*). The hemodynamic response function, ℎ (Stephan et al. 2007b), translates the neural activities into measured BOLD^21^ signals, *y*(*t*), which also contain the observation noise, *e*(*t*). After linearization, Fourier transformation and parametrization of the noise cross-spectra, this generative model admits a deterministic form that can be efficiently *inverted* using variational Bayesian methods (Friston et al. 2007; Friston et al. 2014; Razi et al. 2015).

Model inversion refers to the estimation of optimal model parameters, given a generative model and some observations. In variational Bayesian inference, this is achieved by maximizing free energy (*F*, a lower bound on log model evidence) with respect to the approximate posteriors of the parameters, *Q*(θ|*y*, *m*) (Friston et al. 2007; Zeidman et al. 2022). Notably, free energy offers a trade-off between model accuracy and complexity (F = accuracy – complexity), with the latter term protecting against overfitting. Moreover, maximized free energy serves as a proxy for log model evidence (*F*_*opt*_(*Q*) ≈ log *P*(*y*|*m*)), which is used to compare the plausibility of different models explaining the same data. The associated optimized posteriors (*Q*_*opt*_(θ|*y*, *m*) ≈ *P*(θ|*y*, *m*)) are used to perform inference on the parameters. The most prominent parameters of DCM are the effective connections encoded in the *A*_*n*×*n*_ matrix. Each off-diagonal entry *a*_*ij*_ of this matrix denotes the causal influence of region *j* on region *i*, which can be excitatory (positive) or inhibitory (negative). Conversely, the self-connections^22^ (*a*_*ii*_) are parametrized with negativity constraints to ensure dynamical stability of the model (Zeidman et al. 2019a).

This concludes our brief description of the generative model of spectral DCM and the associated Bayesian model inversion scheme. Having inverted each subject’s DCM, group analysis was conducted using a hierarchical Bayesian framework known as parametric empirical Bayes (PEB)—as elaborated next.

#### 3.4.3 Group analysis using parametric empirical Bayes

PEB is a hierarchical Bayesian model, particularly useful for estimating group effects in DCM studies. Operationally, PEB is a Bayesian general linear model (GLM) that partitions between-subject variability into certain designed group effects (such as group mean and difference) and some additive random effects. In contrast to the *summary statistic* random effects approach based on point estimates, PEB accommodates the *full posterior* densities of the parameters. In its simplest form, PEB can be expressed as:

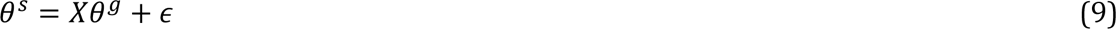

where 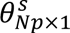 includes the posteriors of *p* DCM parameters for *N* subjects; *X*_*Np*×*pd*_ is the design matrix for *d* regressors replicated over *p* parameters; 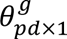 denotes the group parameters; and ϵ_*Np*×1_ contains the random effects. The columns of X encode the hypothesized sources of inter-subject variability (e.g., group mean). To invert a PEB model, subject and group level parameters are estimated iteratively in a variational Bayesian scheme (Friston et al. 2007; Friston et al. 2015; Friston et al. 2016; Zeidman et al. 2019b). That is, group-level parameters are estimated by assimilating the posteriors of subject-level parameters. Then, group parameters act as empirical priors for reinversion of subject-specific DCMs. This iterative procedure continues until convergence. The empirical priors are like *typical* group values, which serve to guide subject-level inferences and help to avoid local maxima problems (Friston et al. 2015; Friston et al. 2016; Zeidman et al. 2019b).

In the present study, PEB was used to compute the group mean value for each effective connectivity parameter. The PEB structure was further optimized post-hoc using exploratory BMR (Friston et al. 2016; Friston and Penny 2011). These PEB (i.e., group DCM) posteriors comprised the directed cyclic graph for which causal centralities of the nodes were computed (using Eq. 6) and compared against some major graph theoretical centralities—as explained next.

### 3.5 Causal vs. graph-theoretical centralities

The PEB posterior encodes the average causal influences among the regions of eDMN. For this cyclic graph, the causal centralities of individual (cortical and subcortical) regions were computed using Eq. 6. Then, several graph-theoretical node centralities were computed on the same network. These included the *strength*, *betweenness*, *closeness* and *eigenvector* centralities, which are succinctly introduced here, and mathematically defined in Appendix C, based on (Bonacich 2007; Bonacich and Lloyd 2001; Freeman 1979; Opsahl et al. 2010).

Briefly, node *strength* computes how strongly a node is directly connected to the other nodes in the network (by summing over the absolute edge weights connected to that node). *Betweenness centrality* quantifies the bridging/flow-passing role of a node in the network (by computing how often it lies on the shortest paths between the other nodes). *Closeness centrality* reflects the efficiency of a node in spreading information over the network (computed as the reciprocal of the average shortest path from that node to all the other nodes). Finally, a node with high e*igenvector* centrality is connected to multiple other highly central nodes (which is formally computed using the first eigenvector of the graph adjacency matrix). From the perspective of *centrality typology*, these measures cover the major three types of centralities defined in (Borgatti and Everett 2006), and the same measures have been studied by (Dablander and Hinne 2019) in the context of causal inference on DAGs.

Generally, more central nodes in a graph contribute further towards the connectedness or the flow of information over the network. So, intuitively, one might expect that these graph-theoretically central nodes possess high causal power over the network as well. The comparative analysis in the next section is meant to demystify this point, through the working example of eDMN’s causal network.

### 3.6 Causal relevance of different centralities

DCM adopts a fully Bayesian approach to parameter estimation, with Gaussian *shrinkage priors* (*P*(θ|*m*)) that constrain the estimates of causal connections (i.e., *A* matrix in Eq. 7). The shrinkage priors have zero mean and small identical prior variances, *N*(μ = 0, Σ = 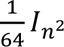), and thus *shrink* the posterior estimates of the causal connections towards zero. Crucially, the higher the prior variance is set, the easier it becomes for the posterior connection estimates to deviate from the prior expectation of zero—if this is sufficiently supported by the observed data. As such, if apriori we believe that certain causal connections are more probable to be nonzero, then incorporating this information during the model fitting procedure (by increasing the prior variances of those connections) will improve the model evidence. This notion has been validated in the context of structurally-informed DCMs, where higher probability of anatomical connections between regions (estimated via probabilistic tractography) was encoded as higher probability of causal connections during DCM inversion, which resulted in significantly improved group model evidence, and confirmed the causal relevance of the anatomical information for models of effective connectivity (Sokolov et al. 2019; Sokolov et al. 2020; Stephan et al. 2009).

In the present work, this method was applied retrospectively; that is, the centrality information (computed from the group DCM graph) was translated into revised *priors* for the causal connections, to reinvert the same DCMs. If reinversion with centrality-informed priors results in nontrivial improvement in model evidence, this would speak to the causal relevance of the centrality pattern. Moreover, different centrality measures can be compared in terms of their causal significance, based on the extent of change in model evidence.

To translate the centrality information into updated priors on the causal connections, the node centrality values were first linearly mapped to the [1,8] interval, and then used to scale the original (default = 1/64) prior variances of the efferent connections of that node:

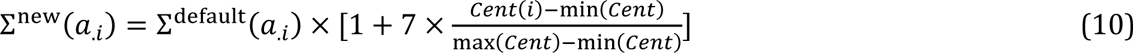

where *Cent*(*i*) refers to the centrality of node *x*_*i*_ and Σ(*a*_.*i*_) denotes the prior variance of the connections originating from node *x*_*i*_. Intuitively, this means that we expect the nodes with higher centrality to exert stronger causal influence over the connected regions. Operationally, model reinversion was implemented using BMR, which has closed-form solutions for the evidence and posteriors of a model with updated priors (Appendix B). Following subject-level DCM reinversion, the updated group-level model evidence was computed using PEB (as explained in section 3.4.3). In brief, this analysis was meant to reveal whether the centrality information would affect the model evidence at the group level.

To examine whether any potential change in model evidence was a mere byproduct of relaxed (i.e., scaled) priors or whether it was indeed specific to the centrality-guided scaling pattern, a control model was set up for each centrality type. In the control models, the centrality scores were randomly shuffled before scaling the prior variances (using Eq. 10) and reinverting the DCMs. The model evidence from the control models would adjudicate on the significance of the original findings.

## 4 Results

### 4.1 Causal structure of the eDMN

Spectral DCM was used to estimate the effective (causal) connections among the regions of eDMN (Table 1), for each subject. For each DCM, model fitting was assessed using the coefficient of determination, *R*^2^, which reflects the proportion of variance in the observations (i.e., cross-spectra) that is explained by the spectral DCM model^23^ (*R*^2^ = 0.91 ± 0.03, across the group). Group analysis was conducted in the PEB framework (section 3.4.3). Group effective connectivity results have been plotted in Fig. 3.

**Fig. 3:**
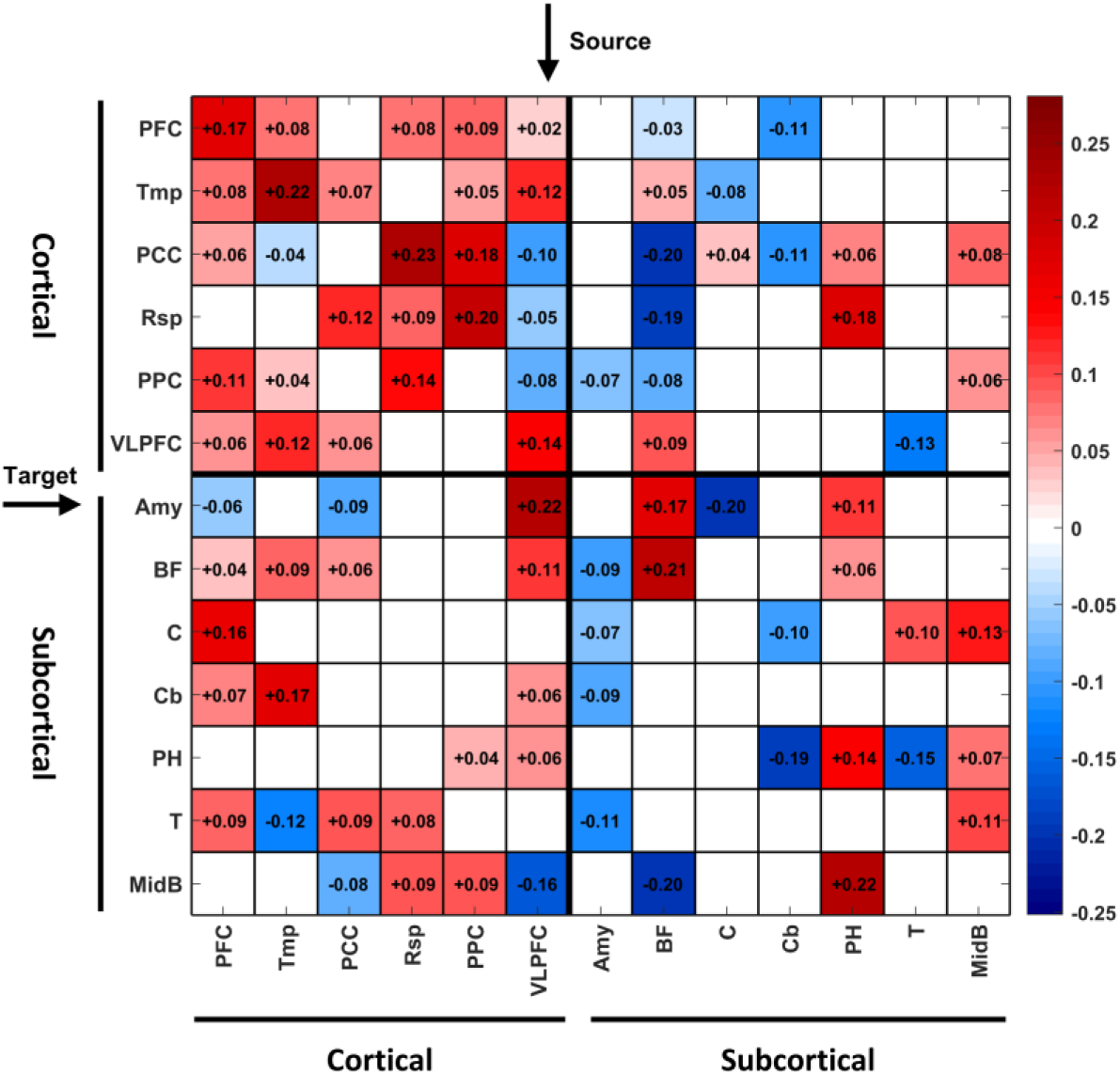
Group DCM results for the eDMN. This figure illustrates the expected PEB posteriors, which specify the average causal architecture of the eDMN in 74 healthy subjects. The columns and rows denote the source and target regions, respectively. The extrinsic and intrinsic/self connections are in units of Hz and log Hz, respectively. Self-connections (on the main diagonal) are always inhibitory, and can be converted to units of Hz using −0.5 ∗ exp (*A*_*ii*_). The posterior uncertainties of the connections are illustrated in Fig S4. Abbreviations for the eDMN regions are available in Table 1.

As for the causal structure of eDMN, Fig. 3 shows that the causal influences among the cortical regions of eDMN are dense and mostly excitatory (apart from some inhibitions induced by VLPFC/Tmp). Conversely, the within-subcortical causal architecture seems sparser and more balanced in terms of excitatory and inhibitory effects. Among the subcorticals, the amygdala and cerebellum act as inhibitors, whereas the parahippocampal region and the midbrain assume excitatory roles. Notably, the cortical-to-subcortical causal influences are mostly excitatory, whereas the subcortical-to-cortical effects are both inhibitory and excitatory (with basal forebrain significantly contributing to both regimes). In Fig. S4 the posterior *uncertainties* of the group effective connections have been illustrated in addition to the posterior expectations. After estimating the average causal influences among the eDMN regions, the node centralities were computed based on this causal graph.

### 4.2 Centrality of the eDMN regions

The group DCM posteriors encode the average causal influences among the regions of eDMN, in 74 healthy subjects. For this graph, the causal and GT centralities were computed for each region, as elaborated in section 3.5. The centrality scores and rankings are included in Table 2. The centrality rankings have also been visualized in Fig. 4-A. In each bar plot, one type of GT centrality ranking has been plotted besides the causal centrality ranking of the regions. Higher ranks correspond to higher centrality values (with the maximum/best rank = 13 = number of regions).

**Fig. 4:**
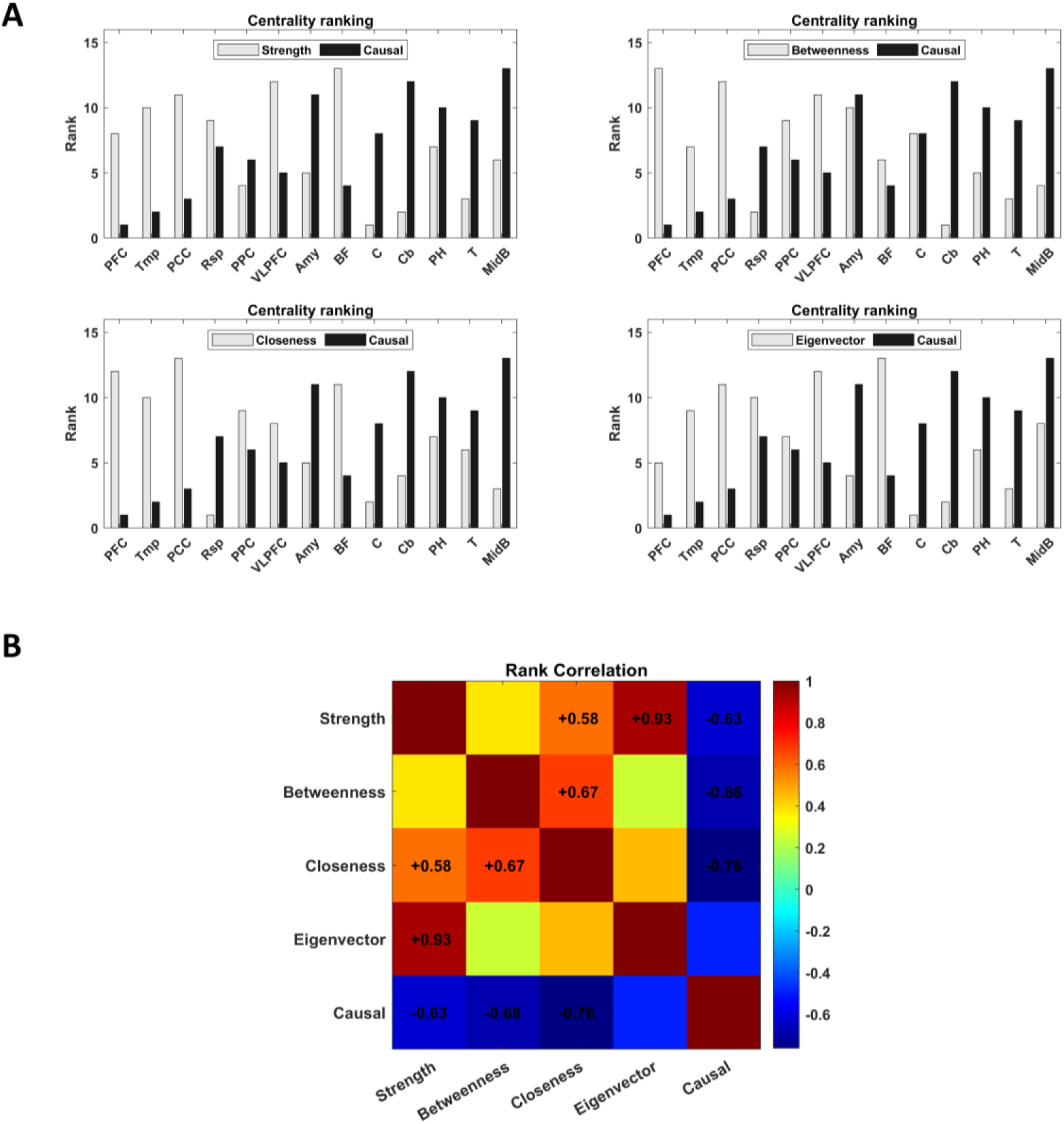
Centrality ranking of the eDMN regions, in the average causal graph (i.e., group DCM graph) of the eDMN. (A) In each bar plot, one type of graph-theoretical centrality ranking (in light gray) has been plotted besides the (black) causal centrality ranking of the eDMN regions. Higher ranks correspond to higher centrality values, which have been listed in Table 2. (B) Rank correlation between the different types of centralities computed on the average eDMN causal graph. Significant correlations (p<0.05) have been annotated. Overall, the graph-theoretical centralities are rather similar in their ranking patterns, while negatively correlated with the causal ranking results.

**Table 2:** Centrality scores and ranks of the eDMN regions, based on different centrality measures applied to the average causal graph (i.e., group DCM graph) of the eDMN. Higher ranks correspond to higher centrality values. The maximum/best rank is 13, which is the number of modeled regions, as listed in Table 1.

|  | Centrality |  |  |  |  |  |  |  |  |  |
| --- | --- | --- | --- | --- | --- | --- | --- | --- | --- | --- |
|  | Strength |  | Betweenness |  | Closeness |  | Eigenvector |  | Causal |  |
| Region | Value | Rank | Value | Rank | Value | Rank | Value | Rank | Value | Rank |
| PFC | 1.45 | 8 | 33 | 13 | 0.88 | 12 | 0.07 | 5 | 2626.35 | 1 |
| Tmp | 1.54 | 10 | 6 | 7 | 0.77 | 10 | 0.09 | 9 | 3581.82 | 2 |
| PCC | 1.67 | 11 | 29 | 12 | 0.99 | 13 | 0.10 | 11 | 6321.73 | 3 |
| Rsp | 1.53 | 9 | 1 | 2 | 0.53 | 1 | 0.09 | 10 | 9708.33 | 7 |
| PPC | 1.24 | 4 | 7 | 9 | 0.72 | 9 | 0.07 | 7 | 9305.69 | 6 |
| VLPFC | 1.73 | 12 | 24 | 11 | 0.72 | 8 | 0.10 | 12 | 7878.70 | 5 |
| Amy | 1.27 | 5 | 7 | 10 | 0.66 | 5 | 0.07 | 4 | 12085.05 | 11 |
| BF | 1.89 | 13 | 5 | 6 | 0.84 | 11 | 0.12 | 13 | 6361.08 | 4 |
| C | 0.87 | 1 | 6 | 8 | 0.58 | 2 | 0.04 | 1 | 10745.96 | 8 |
| Cb | 0.89 | 2 | 0 | 1 | 0.63 | 4 | 0.05 | 2 | 12282.63 | 12 |
| PH | 1.45 | 7 | 2 | 5 | 0.71 | 7 | 0.07 | 6 | 11663.28 | 10 |
| T | 0.98 | 3 | 1 | 3 | 0.68 | 6 | 0.05 | 3 | 10930.42 | 9 |
| MidB | 1.30 | 6 | 1 | 4 | 0.60 | 3 | 0.07 | 8 | 16030.32 | 13 |

It is apparent from Fig. 4-A that the GT rankings are quite distinct from the causal ranking. Specifically, the GT measures assign the higher centrality ranks to the cortical areas, whereas causal centrality attributes the higher ranks to the subcortical regions. For instance, while PCC is highly central in a graph-theoretical sense (i.e., always in the top three basket), it has the 3^rd^ lowest causal centrality rank. Another example is PFC, which has a relatively high rank based on GT measures (average rank = 9.5); however, PFC gets the lowest rank (=1) based on causal centrality. Conversely, the midbrain, which is on top of the causal centrality ranking (rank =13), has a mediocre GT rank of 5.25 on average. Or, the causally important cerebellum (with causal rank = 12), has a very low average GT rank of 2.25.

On average, the GT centrality ranks are 8.96 and 5.32, for cortical and subcortical regions, respectively; however, the opposite relationship holds for the causal centrality ranks: average cortical rank = 4; average subcortical rank = 9.57. In Fig. 4-B, the relationship between the rankings of different centrality measures has been formalized using Spearman’s rank correlation coefficient. The overall trend speaks to a relative consensus among the GT centrality rankings, which stand in striking contrast to the causal centrality results—as reflected by the negative correlations in the last row. The causal significance of these centrality measures were assessed next, by incorporating them as prior information in the causal models.

### 4.3 Causal relevance of different centralities

Once the node centrality scores were linearly mapped to the [1,8] interval (as shown in Fig. 5), they were used to scale the prior variances on the efferent connections of the associated nodes. The DCMs were then reinverted with these updated priors (using BMR), and group analysis was conducted in the PEB framework. The improvement in model evidence (under centrality-informed priors) was recorded at the subject and group level. The control models were set up similarly, but using randomly permuted centralities. The results have been illustrated in Fig. 6.

**Fig. 5:**
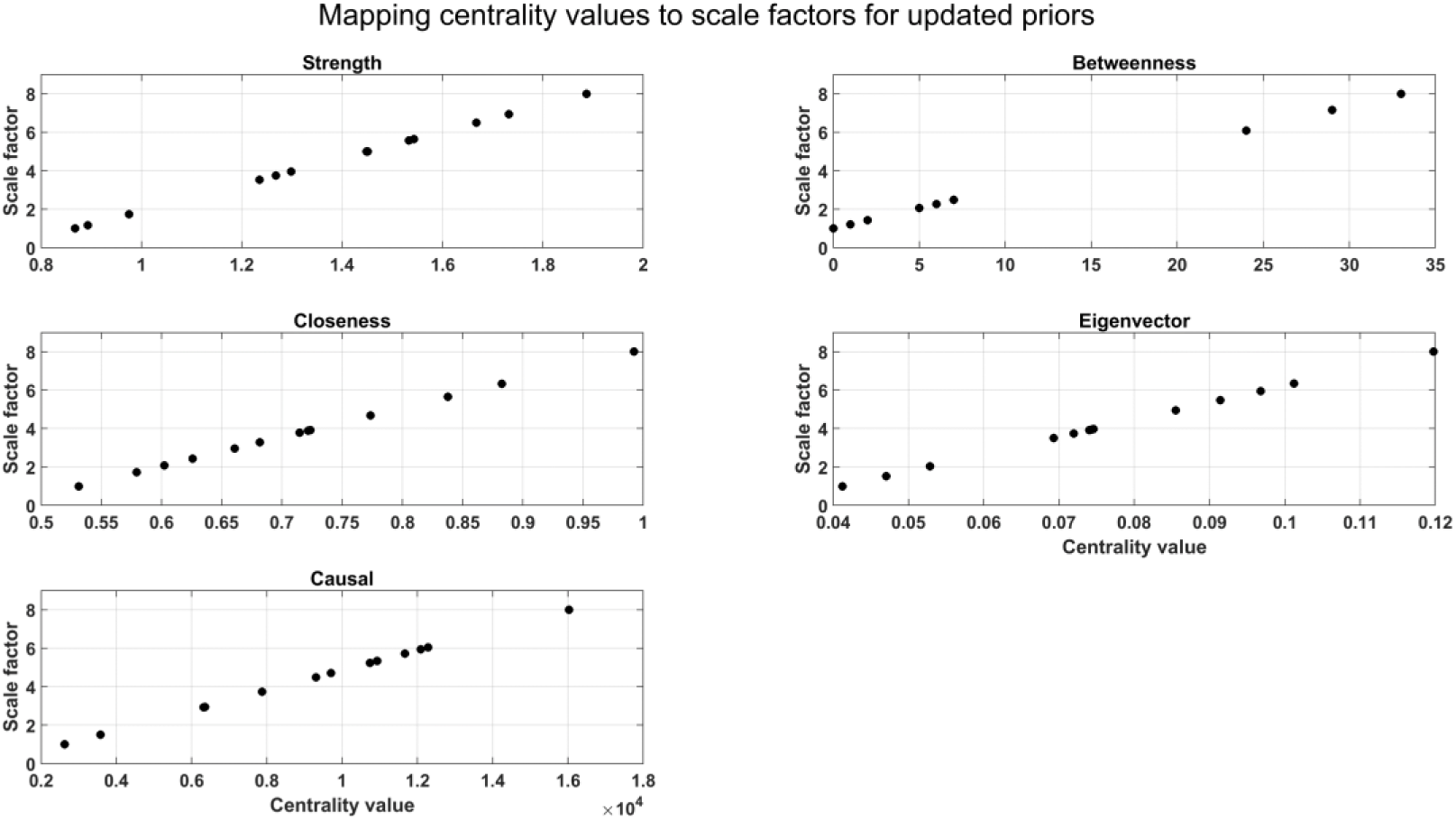
Mapping the node centrality values to scale factors in the [1,8] interval. The mapped values were used to scale the default prior variances on the efferent connections of the associated nodes, based on Eq. 10.

**Fig. 6:**
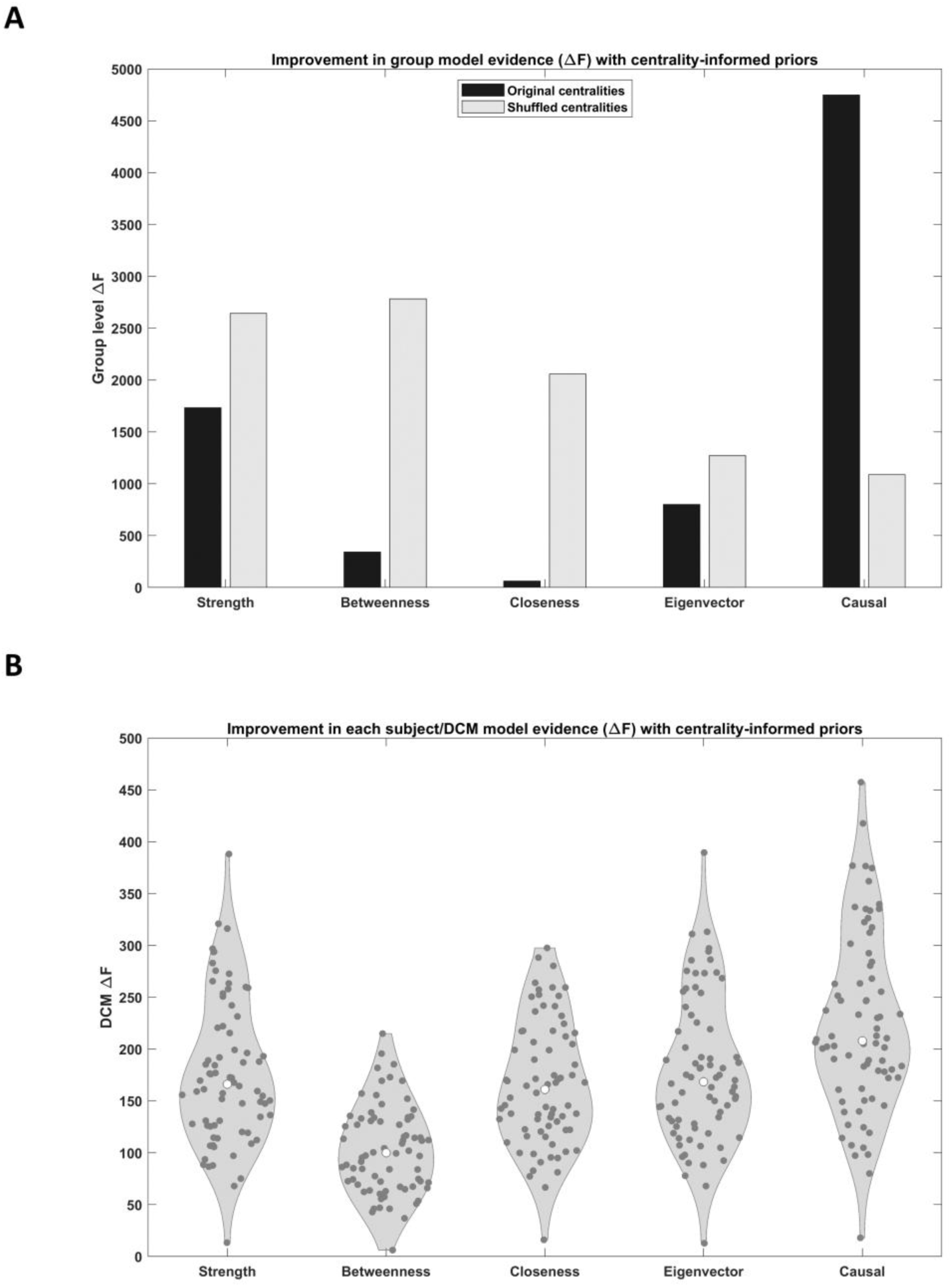
Causal relevance of different centralities. (A) Improvement in group model evidence, following DCM reinversion under centrality-informed priors (black bars), and for control models with shuffled centralities (light gray bars). (B) Improvement in subject model evidence, following DCM reinversion under centrality-informed priors. Each dark circle in the violin plots corresponds to Δ*F* for one subject/DCM. Median values are marked with white circles.

At first glance, all the centrality-informed models seem to have improved the model evidence per subject and over the group (Fig. 6). However, for the GT centralities, this improvement is a simple byproduct of relaxing/scaling the priors—not the specific centrality pattern used to scale the priors. This can be inferred from Fig. 6-A, which shows that the control models (with shuffled centralities) have achieved higher improvement in model evidence than the original centrality-informed models. Only when the causal centrality scores were encoded as priors did the improvement in model evidence become meaningful—by significantly outperforming the control model (Δ*F* = Δ*F*_*orig*_ − Δ*F*_*control*_ = 3659.3). Note that, in the spirit of log Bayes factors^24^, |Δ*F*| > 3 nats is interpreted as “strong evidence” in favour of one model against another, while |Δ*F*| > 5 nats denotes “very strong evidence” (Kass and Raftery 1995).

Also note that, the average subject-level improvements in model evidence (Fig. 6-B) follow the same trend as the group-level improvements (Fig. 6-A, black bars); except when comparing the closeness and betweenness centralities. That is, while Δ*F* is on average higher per subject/DCM for the closeness (than betweenness) centrality, the group Δ*F* is lower for the closeness (than betweenness) centrality-informed model. This is because, the group-level Δ*F* of PEB is not merely the sum of subject-level Δ*F*’s—unlike a fixed effects approach to group Δ*F* computation (Stephan et al. 2007b; Stephan et al. 2007a; Summerfield and Koechlin 2008). Instead, by virtue of accounting for the between-subject (i.e., random) effects, the group-level free energy of PEB is the expected sum of subject-level free energies (under the approximate posterior of the group-level parameters) minus the group-level complexity term^25^, as shown in (Friston et al. 2015; Friston et al. 2016). The latter term can explain the disparity between the subject and group level change in model evidence, when comparing the closeness and betweenness centrality-informed models. This highlights the importance of the Bayesian hierarchy of PEB, which explicitly models the between-subject variations (i.e., random effects) in the causal networks. As such, the group-level results are more complete and reliable.

## 5 Discussion

When it comes to assessing *centrality* in dynamic causal models, graph-theoretical measures fall short. This is because, topology-based GT centralities do not account for the *dynamics* of the specific generative model behind the graph (Borgatti 2005) and they are not qualified to deliver *causal* verdicts either (Dablander and Hinne 2019). Hence, a new *causal centrality* measure was introduced in this paper, based on the notion of *intervention* on the edges of a graphical model. It was shown that this measure is *identifiable* by means of Bayesian model reduction. As a proof of concept - and to compare the new measure with major GT centralities - a worked example was presented using the dynamic causal model of the extended default mode network (Alves et al. 2019). After estimating the average DCM graph of the eDMN (from 74 healthy subjects), causal centrality was computed for each region, and compared against common GT centralities. Moreover, the causal relevance of different centralities were assessed in a model comparison framework, where the centrality information modulated the prior (variance) of the causal connections in the DCM—and the ensuing change in model evidence reflected the causal relevance of the centrality information. In the following, the key aspects of this study and their implications are discussed.

Causal centrality was defined as the distributional divergence induced by intervention on the edges of the causal graph (Janzing et al. 2013; Pearl 2010). This interventional approach is conceptually similar to “lesioning” (Deco et al. 2017; Irimia and van Horn 2014), “hub-opathy” (Bell and Shine 2016) and “vulnerability analysis” (Iturria-Medina et al. 2008; Rawls et al. 2022) in network (neuro)science, which measure the effect of edge/node removal on the network integrity and performance (Gol’dshtein et al. 2004). Notably, since causal centrality is conditioned on the generative model of the DCM associated with the graph, it explicitly accounts for the network *dynamics*. This is important because, classical centrality measures rely only on network *topology*, while the role of individual elements in collective behavior depends inevitably on the specificities of the network *dynamics* (Borgatti 2005; Borgatti and Everett 2006; Bringmann et al. 2019; Klemm et al. 2012; Li et al. 2012).

For instance, in an epidemic spreading model with given topology, picking the GT centrality that can best identify the influential spreaders depends critically on the infection rate— which is a dynamical parameter (Liu et al. 2016). As such, a “dynamics-sensitive” centrality measure was proposed by (Liu et al. 2016) for epidemic processes, which integrates topology and (epidemiological) dynamics. Other “dynamical influence” measures have also been proposed in different contexts, based on spectral decomposition of the Laplacian or adjacency matrix (Klemm et al. 2012; Masuda and Kori 2010; Restrepo et al. 2006) and information flow on undirected graphs (van Elteren et al. 2022). In neuroscience, (Deco et al. 2017) used computational modeling and sequential node removal to show (using an information-theoretic measure) that the critical regions for “information encoding and integration” in the brain are not necessarily the “highest-degree” or “rich-club” members from GT analysis of the *structural connectome*. The present work extends their findings, and argues that GT analysis on the *effective connectome* is not indicative of the dynamical (causal) influence of the regions either—and that dynamics-sensitive measures like causal centrality are needed for this purpose.

Operationally, causal centrality was computed using Bayesian model reduction. BMR can analytically compute the approximate model evidence and posteriors of a submodel, from the same quantities (and the priors) already available for the full model. A submodel (aka a reduced model) includes a subset of the original model’s parameters, with the rest of the parameters fixed to zero (using precise null priors). Hence, given the priors and posteriors of the full model, BMR can answer the question: what would the posteriors have been, had the priors assumed a reduced form?^26^ The answer that BMR estimates would render the post-intervention distribution (in Eq. 4) *identifiable*. As such, even though the main function of BMR has so far been in post-hoc model optimization and structure learning (Beckers et al. 2022; Friston et al. 2016; Friston et al. 2018; Friston and Penny 2011; Jafarian et al. 2019; Neacsu et al. 2022), another useful application is in the computation of causal centrality.

We now review the results from dynamic causal modeling the extended default mode network. In the causal architecture of the eDMN (Fig. 3), cortical regions had dense and (mostly reciprocal) excitatory connections among themselves, except for some inhibitions induced by the VLPFC and temporal regions. Conversely, the within-subcortical effective connections were sparser and rather balanced in terms of excitation and inhibition. In this causal model, the amygdala and cerebellum exerted inhibitory influences on the other (subcortical and cortical) regions, whereas the parahippocampal and midbrain regions had excitatory roles. The basal forebrain exerted a mixture of excitatory and inhibitory effects, and it was the only subcortical structure of this causal model that affected all the cortical regions of the eDMN—mostly as an inhibitor. Overall, the subcortical-to-cortical effects were both inhibitory and excitatory, whereas the cortical-to-subcortical influences were predominantly excitatory. The latter is reminiscent of the presumed “funneling” (i.e., information integration) role of the subcortical nuclei (Bell and Shine 2016; Parent and Hazrati 1995), while the former could echo the subcortical “neuromodulatory” effects (Avery and Krichmar 2017; Robbins and Arnsten 2009).

Subsequently, based on the average causal graph (from DCM group analysis), the eDMN regions were assessed in terms of their causal and GT centralities. Overall, the *subcortical* regions turned out to be more causally central, whereas the *cortical* regions were more graph-theoretically central (Fig. 4-A). Specifically, the midbrain achieved the highest causal centrality, followed by the cerebellum, amygdala, parahippocampal region, thalamus and caudate. The only subcortical structure with low causal centrality (and high GT centrality) was the basal forebrain. The basal forebrain has high degree and betweenness centrality in the *structural* connectome of the eDMN, as reported in (Alves et al. 2019). Herein, the basal forebrain achieved the highest strength and eigenvector centralities in the effective connectome (i.e., the causal graph) of the eDMN (Fig. 4-A), but in terms of causal centrality it occupied a low rank (of 4). This is an example of divergent GT and causal centrality judgements based on the same causal graph. Another example is the thalamus, which is reportedly highly central in the *structural* connectome of the eDMN (Alves et al. 2019), and causally central in the effective connectome as well (rank = 9), whereas graph theory assigned a low centrality value to the thalamus (average rank = 3.75) in the causal graph.

Overall, the four GT centrality rankings were more consistent with each other, whereas they were negatively correlated (or uncorrelated) with the causal centrality ranks (Fig. 4-B). This is important because, causal centrality accounts for the dynamics of the (biophysical) generative model of DCM, whereas the graph-theoretical measures are either dynamics-free or uninformed of the specific dynamics on the graph (Borgatti 2005; Borgatti and Everett 2006). As a result, only the (dynamics-aware) causal centrality detected the influential role of the subcortical regions in the extended DMN (Fig. 4).

There is accumulating evidence about the contribution of subcortical structures to the integration of large-scale networks (Bell and Shine 2016). Specifically, the implication of subcortical regions in the organization of the DMN has been acknowledged in recent structural (Alves et al. 2019; Bzdok et al. 2013; Cunningham et al. 2017), functional (Alves et al. 2019; Bär et al. 2016; Buckner et al. 2011; Bzdok et al. 2013; Choi et al. 2012; Cunningham et al. 2017; Di Martino et al. 2008; Fransson 2005; La Cruz et al. 2021; Lee and Xue 2018; Li et al. 2021; Roy et al. 2009; Stoodley and Schmahmann 2009) and effective connectivity (Harrison et al. 2022; La Cruz et al. 2021; Nair et al. 2018) analyses, as well as electromagnetic (Elias et al. 2021; Gratwicke et al. 2013; Kakusa et al. 2020; Schiff et al. 2007), ultrasonic (Cain et al. 2021a; Cain et al. 2021b; Monti et al. 2016) and optogenetic (Klaassen et al. 2021; Lozano-Montes et al. 2020) stimulation studies, and pharmacological experiments (Carhart-Harris et al. 2013; Kelly et al. 2009; Kunisato et al. 2011; Metzger et al. 2016; van de Ven et al. 2013; van Wingen et al. 2014).

Subcortical structures contain rich neurochemical nuclei, and large receptive and projective fields. As such, the neuromodulation of small subcortical nuclei can affect large and distributed portions of the cortex (Horn et al. 2017; Horn et al. 2019; Horn and Fox 2020; Li et al. 2021; Schiff et al. 2007). Importantly, classical GT centrality measures seem unable to reflect this critical subcortical functionality. For instance, in the graph-theoretical study of (Bär et al. 2016), the dopaminergic midbrain nuclei were not identified as central nodes within the DMN, whereas the VMPFC and PCC showed up as the usual “key-hubs”. This is in line with the graph-theoretical results of the present study (Fig. 4-A), where the PCC, VLPFC and PFC achieved rather high GT centralities, while the subcortical regions received the lower values. However, when the network dynamics were accounted for, the midbrain achieved the highest causal centrality rank, followed by the other subcortical structures— and the cortical regions moved down the list. Importantly, *only* the causal centrality pattern improved the model evidence nontrivially in the model comparison framework (Fig. 6).

The model comparison revealed that: if the prior probabilities of the causal influences are increased in proportion to the causal centralities of their source regions, then the DCM model evidence would improve *significantly*—i.e., beyond the mere effect of relaxed priors. However, for the GT centralities, the improvement in model evidence was insignificant; i.e., the centrality-informed models were outperformed by the control models (with randomly permuted centralities). As such, the only causally-relevant measure turned out to be causal centrality, which happened to be the sole centrality that endorsed the prominent role of subcortical structures in the causal architecture of the eDMN.

Notably, many of the subcortical structures – which contain rich neurochemical nuclei - are structurally and functionally disrupted in various neurological and psychiatric disorders (Bocchetta et al. 2021; Dandash et al. 2014; Jauhar et al. 2018; Kirschner et al. 2022; Koshiyama et al. 2018; Lecciso and Colombo 2019; Lorenzini et al. 2021; Panda et al. 2022; Park et al. 2019; Sabaroedin et al. 2023a; Sabaroedin et al. 2023b; Výtvarová et al. 2017; Wolf et al. 2011; Yamamoto et al. 2022; Zeng et al. 2023; Zhao et al. 2022). Specifically, neurodegenerative disorders that are characterized by early and selective subcortical pathology − such as Parkinson’s disease and Huntington’s disease (Braak et al. 2003; Vonsattel et al. 1985) − are also associated with fragmentation of the global network topology in early-stage disease (Harrington et al. 2015; Luo et al. 2015; McColgan et al. 2015; Olde Dubbelink et al. 2014; Sang et al. 2015), which deteriorates with disease progression (Harrington et al. 2015; McColgan et al. 2015; Olde Dubbelink et al. 2014). Such indirect evidence for the “integrative” role of subcortical structures (Bell and Shine 2016) corroborates the neuroanatomical evidence about their large projective and receptive fields, which has led to their inclusion as potential *therapeutic intervention* targets (Alosaimi et al. 2022; Georgiev et al. 2021; Horn et al. 2017; Horn et al. 2019; Horn and Fox 2020; Schiff et al. 2007).

Unraveling the *mechanisms* of different clinical interventions, and *designing* suitable stimulation protocols are trending topics in computational and clinical neuroscience, which draw upon methods from *perturbation analysis* (Deco et al. 2019; Mana et al. 2022; Sanz Perl et al. 2022; Vohryzek et al. 2022) and *control theory* (Gu et al. 2017; Gu et al. 2022; Kamiya et al. 2023; Srivastava et al. 2020; Yang et al. 2021). Despite technical variations, the conceptual procedure remains the same: model the brain as a dynamical system; fit this model to the spatiotemporal features of the brain in the unperturbed state; add a stimulation term to the model (or modulate some model parameters) to trigger transition to a new brain state; finally, optimize the location and profile of the stimulation or modulation for (in silico) transition to the new state. Depending on the study, the source/target brain *states* may correspond to the *profiles of transient brain dynamics* in normal/pathological (Mana et al. 2022; Vohryzek et al. 2022) or sleep/awake (Deco et al. 2019) scenarios; alternatively, the *states* may be defined as the *spatiotemporal patterns* of rest/task (Kamiya et al. 2023) or rest/stimulation (Yang et al. 2021) conditions.

Notably, optimizing the stimulation locations(s) is usually performed using exhaustive search, which is computationally expensive (Deco et al. 2019; Mana et al. 2022; Vohryzek et al. 2022). Recent work suggests that the nodes’ *controllability^27^* at rest can help to identify the optimal stimulation locations (Yang et al. 2021). Since average controllability of a node is strongly correlated with its *causal outflow^28^* (Cai et al. 2021), and network controllability mediates the *dynamical flexibility^29^* of the brain to transition between states (Gu et al. 2022), one might speculate that causal measures of centrality may be used as *proxies* for controllability metrics, to constrain the search space for optimal intervention sites that can induce specific state transitions. The advantage of using causal centrality - as a proxy for controllability - would be in the computational efficiency of the former by virtue of Bayesian model reduction. Further research is required to formalize the relationship between causal centrality and controllability metrics in neuronal networks, and to study the implications for identifying therapeutic stimulation targets (Alosaimi et al. 2022; Eraifej et al. 2023; Ezzyat et al. 2018; Georgiev et al. 2021; Wang et al. 2022; Zangen et al. 2023).

The more immediate application of causal centrality would be to elucidate the influence of different regions in the causal models of neurotypical functional networks (such as the eDMN (Alves et al. 2019) and attention network (Alves et al. 2022)), during resting state and task conditions. A follow-up application would be to compare the neurotypical (causal centrality) results with those obtained in pathological conditions, e.g., in schizophrenia (Zarghami et al. 2020; Zarghami et al. 2023). In a recent study, (Mana et al. 2022) used perturbation analysis to identify a set of “critical” regions whose modulation can push a healthy brain towards the pathological dynamics of schizophrenia. Comparing these critical regions with the causally central hubs in the healthy and schizophrenic brain would be mechanistically insightful. Especially if applied to early-onset unmedicated patients (Anticevic et al. 2015), alterations in causal centrality might shed light on the aetiology of the disorder. Moreover, treatment response to different antipsychotic medications, which has previously been studied based on dynamics-free GT measures (Hadley et al. 2016), could be analyzed from the perspective of dynamics-sensitive causal centrality alterations.

As for the limitation of the current work, note that although the dynamics *on* the network were accounted for by the DCM, the dynamics *of* the network (i.e., the temporal evolution of the graph structure (Bassett and Sporns 2017)) were not modeled here. As such, the estimated causal centralities represent session-average values. More advanced DCMs can model the dynamics *of* the network as well, in continuous or discrete time (Jafarian et al. 2021; Zarghami and Friston 2020). In future work, these models can be combined with the formalism of *multilayer networks* (Huang and Yu 2017; Kivela et al. 2014) in applied mathematics to derive causal centrality measures for evolving causal networks. Moreover, even though the present paper dealt with *nodal* causal centrality, it is perfectly possible to quantify *edge* causal centrality (Faskowitz et al. 2020; Novelli and Razi 2022; Zamani Esfahlani et al. 2020) using the same interventional approach.

Finally, although dynamic causal modeling is best known in the field of neuroimaging where it was first introduced (Friston et al. 2003; Moran et al. 2013), the overall framework (which consists of generative modeling of coupled dynamical systems plus Bayesian model inversion) and the computational devices that come along with it (such as hypothesis testing, uncertainty quantification, Bayesian model reduction, Bayesian group analysis, etc.) are quite generic. For instance, a recent epidemiological DCM has been developed to model viral spread among geographical regions (Friston et al. 2020). As such, causal centrality can be applied to different sorts of DCMs—in neuroimaging and other fields—to identify the key players in the collective dynamics of the causal model.

## 6 Conclusions

This paper introduced causal centrality, a dynamics-sensitive measure for assessing the causal importance of nodes/edges in a class of time-dependent cyclic graphs, known as dynamic causal models. The measure was defined as the normalized KL-divergence between the pre- and post-intervention distributions of the causal model, which simplified to an identifiable expression using Bayesian model reduction. Face and construct validation was established by dynamic causal modeling the extended DMN (in 74 healthy subjects), following which causal centralities of the regions were computed for the group-average causal graph, and contrasted against major graph-theoretical centralities. As such, the subcortical regions of the eDMN turned out to be highly causally central, even though the (dynamics-free) graph-theoretical centralities downplayed their roles. Notably, model comparison revealed that only the pattern of causal centrality was causally relevant. These results are consistent with the crucial role of the subcortical structures in the neuromodulatory systems of the brain, and highlight their implication in the organization of large-scale networks. Finally, the potential applications of this new measure - for studying neurotypical and pathological functional networks - were discussed. Future work can elucidate the relationship between causal centrality and controllability metrics, with the prospect of effective target selection for intervention in the brain networks. Causal centrality can also be applied to DCMs developed for applications other than neuroimaging.

## Software note

A demo code for the computation of causal centrality is available at: https://github.com/tszarghami/CausalCentrality. SC-ICA is part of the Group ICA of fMRI Toolbox (GIFT): http://trendscenter.org/software/gift/. Spectral DCM, PEB and BMR have been implemented in SPM12: https://www.fil.ion.ucl.ac.uk/spm/. Graph-theoretical measures can be computed using the *centrality* function in MATLAB.

## Data availability

The data analyzed in this study were obtained from the COllaborative Informatics and Neuroimaging Suite Data Exchange tool (COINS; https://coins.trendscenter.org). Data collection was performed at the Mind Research Network, and funded by a Center of Biomedical Research Excellence (COBRE) grant 5P20RR021938/P20GM103472 from the NIH to Dr. Vince Calhoun. The improved parcellation map of the DMN (Alves et al. 2019) has been publicly shared by the authors on the NeuroVault repository: https://identifiers.org/neurovault.image:568084.

## Author contributions

TSZ: Conceptualization, Methodology, Analysis, Interpretation, Writing.

## Competing interests

The author has no relevant financial or non-financial interests to disclose.

## Appendix A- Causal centrality derivation

In this appendix, the identifiable expression in Eq. 6 is derived from the definition of causal centrality in Eq. 4, and the normalization constant (Z in Eq. 4) is clarified. To start, we expand the KL-divergence between the pre- and post-intervention distributions:

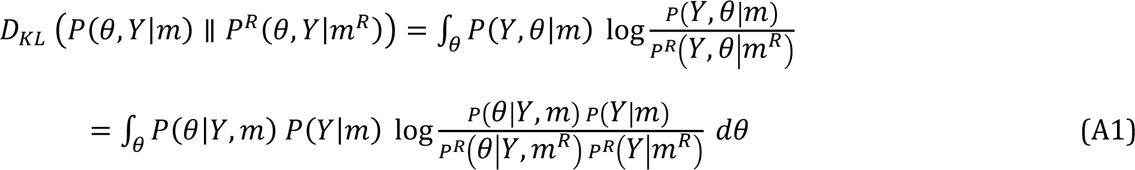

The first equality is just the definition of KL-divergence: 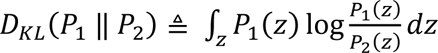.

The second equality uses the product rule of probability: *P*(*Y*, θ|*m*) = *P*(θ|*Y*, *m*) *P*(*Y*|*m*). Herein, R as a superscript refers to the reduced/post-intervention model, whereas the full/pre-intervention model bears no superscript. With some re-arrangement, Eq. A1 simplifies to:

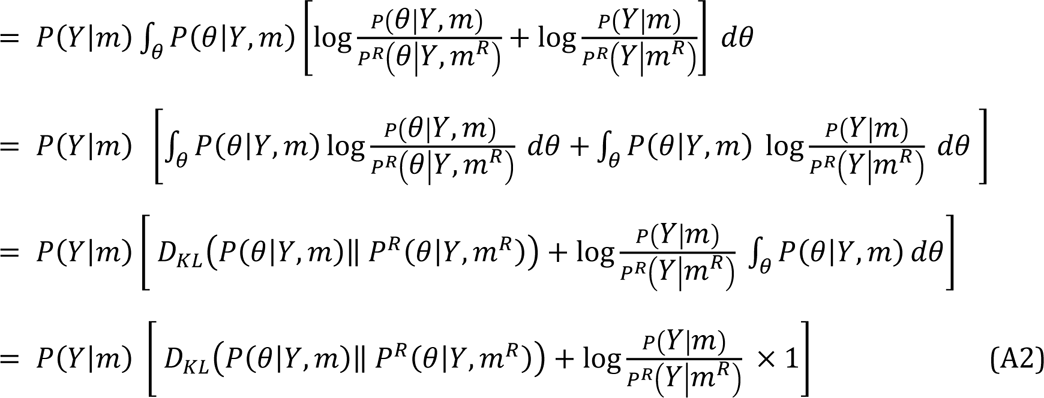

In variational Bayesian inference, the optimized variational density and free energy replace the posterior density and log model evidence: *Q*(θ|*Y*, *m*) ≈ *P*(θ|*Y*, *m*) and *F* ≈ log *P*(*Y*|*m*). Hence, Eq. A2 simplifies to:

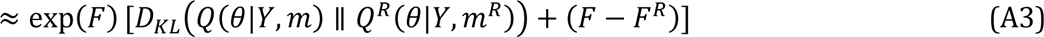

Causal centrality is estimable as the normalized version of Eq. A3, where the normalization constant *Z* = *P*(*Y*|*m*) ≈ exp(*F*). As such, we arrive at Eq. 6:

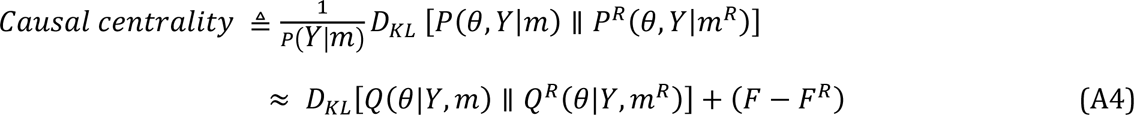

where the posterior and free energy of the reduced model can be computed analytically using Bayesian model reduction, as elaborated in Appendix B. Moreover, under Gaussian assumptions on the posterior distributions in variational Laplace (Friston et al. 2007; Zeidman et al. 2022), the KL-divergence between the two (full and reduced) Gaussian posteriors can be readily computed as:

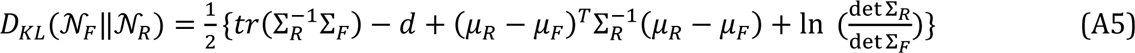

where μ and Σ denote the mean vector and the covariance matrix, respectively; *d* is the dimensionality of the multivariate Gaussians; *F* and *R* subscripts refer to the full and reduced models, respectively; *tr* stands for the trace of a matrix, and ln = log_*e*_.

## Appendix B- Bayesian model reduction derivations

Bayesian model reduction refers to the analytic inversion of reduced models using the priors and posteriors of a full model. Reduced models are nested within a full model; that is, they include only a subset of the parameters of the full model after “switching off” the other parameters (by imposing very precise null priors, which shrinks them to zero). Consider Bayes rule replicated for the full and reduced models (denoted by *m* and *m*^*R*^, respectively) (Friston et al. 2016; Friston et al. 2018; Friston and Penny 2011):

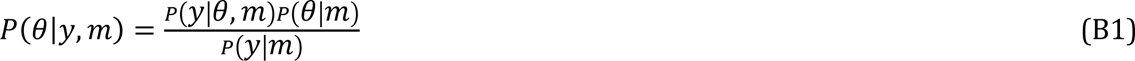

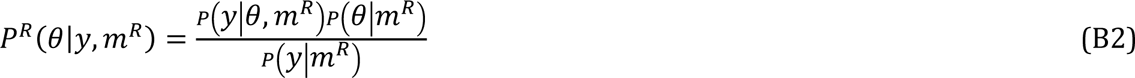

Since the models differ only in terms of their priors, the likelihood terms are identical:

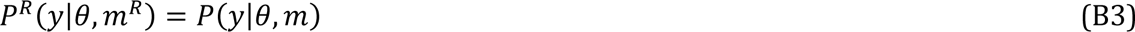

Hence, equating the expressions in Eq. B1 and Eq. B2 over the likelihood gives:

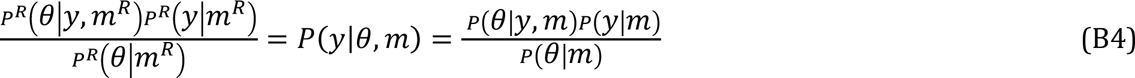

By re-arranging Eq. B4, we get the posterior distribution over the parameters of the reduced model:

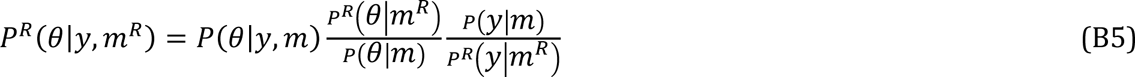

On the R.H.S. of Eq. B5 only *P*^*R*^(*y*|*m*^*R*^) is unknown, which is the model evidence for the reduced model. This term can be obtained by integrating both sides of Eq. B5 over θ:

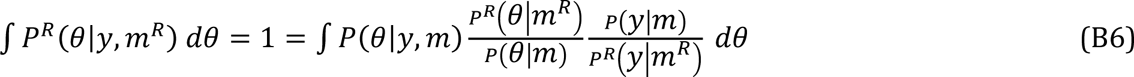

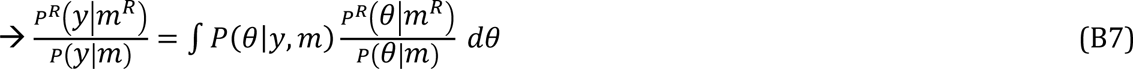

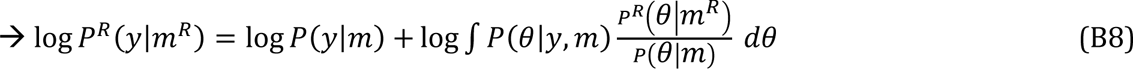

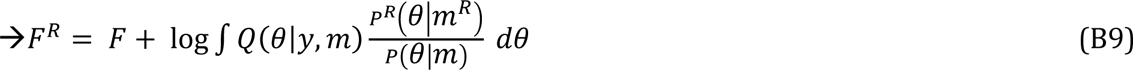

In the last line, the optimized variational density and free energy have replaced the posterior density and log model evidence: *Q*(θ|*Y*, *m*) ≈ *P*(θ|*Y*, *m*) and *F* ≈ log *P*(*Y*|*m*). Under Gaussian assumptions for the distributions (in variational Laplace), the reduced posterior and free energy take simple forms, as elaborated in (Friston et al. 2016; Friston et al. 2018; Friston and Penny 2011).

## Appendix C- Graph theoretical centralities

This appendix includes the mathematical expressions for several graph-theoretical centrality measures. These measures are defined for weighted networks, based on (Opsahl et al. 2010), which generalizes the definitions originally proposed for binary networks (Freeman 1979).

*Node strength* is the generalization of node degree (i.e., the number of connections of a node) for weighted networks, which is defined for node *x*_*i*_ as:

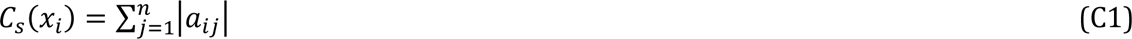

where *n* is the number of nodes, |.| denotes the absolute value function, and *a*_*ij*_ is the (*i*, *j*)^*t*ℎ^ entry of the weighted adjacency matrix *A*.

*Closeness centrality* is defined as:

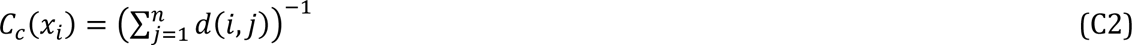

where *d*(*i*, *j*) denotes the shortest path between nodes *x*_*i*_ and *x*_*j*_, which is the path that minimizes the cost of travelling from *x*_*i*_ to *x*_*j*_; the cost of travelling is the inverse of the weight of the (directed) edges connecting the two nodes—as encoded in the adjacency matrix.

*Betweenness centrality* is defined as:

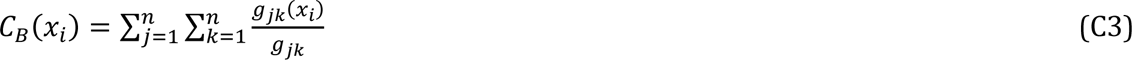

where *g*_*jk*_(*x*_*i*_) is the number of shortest paths between nodes *x*_*j*_ and *x*_*k*_ that go through node *x*_*i*_; and *g*_*jk*_ is the total number of shortest paths between nodes *x*_*j*_ and *x*_*k*_.

Note that in the formulation of DCM (Eq. 7), *A* encodes the directed causal influences; hence, it corresponds to the adjacency matrix of a (directed weighted) *signed* graph, in which excitatory (positive) and inhibitory (negative) effects are equally important. So, to compute the *closeness* and *betweenness* centralities (based on the notion of shortest paths), *A* was converted to an *unsigned* matrix first, using the absolute function: *A* → |*A*|.

Finally, for a node to have high *eigenvector centrality*, it should be connected to many other nodes that also have high eigenvector centrality. In other words, connection to an important node counts more than connection to a less important node. This notion is formalized using the first eigenvector of the adjacency matrix (Bonacich 2007; Bonacich and Lloyd 2001):

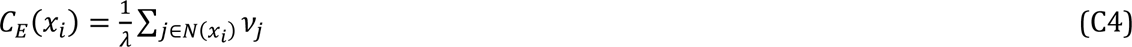

where *λ* is the largest eigenvalue of an undirected adjacency matrix, and *v*_*j*_ is the *j*^*t*ℎ^ element of the corresponding eigenvector; *N*(*x*_*i*_) denotes the set of nodes that are adjacent/neighbor to *x*_*i*_. Since eigenvector centrality is defined for *undirected* graphs, the (unsigned weighted) *directed* adjacency matrix |*A*| was first symmetrized as follows: 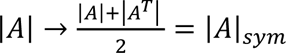; where *T* denotes the transpose operator.

The graph-theoretical centrality measures were computed using the *centrality* function in MATLAB R2021b.

## Footnotes

1 A graph consists of nodes and edges, which represent the network elements and their interrelations, respectively (Barabási (2014)).

2 More precisely, after the *distributions* over the causal strengths have been estimated.

3 Generally, θ encompasses all the unobservable variables of the model, including the hidden states, parameters and hyperparameters. In this work, we only intervene on the causal influences (i.e., the effective connectivity parameters) of DCM, which shape the causal architecture of the graph.

4 Free energy is a lower bound on log model evidence, hence known as evidence lower bound (ELBO) in machine learning. In variational Bayesian inference, the optimized free energy serves as a proxy for log model evidence. Note that this variational free energy is the negative of free energy in statistical physics.

5 Here we make sure that the full model has been structurally optimized (using exploratory BMR); hence, intervention *decreases* the model evidence. Note that, for an over-parametrized model, edge (parameter) removal can *increase* the model evidence, which is the foundation of structure learning and optimization using Bayesian model reduction (Beckers et al. (2022); Friston and Penny (2011); Jafarian et al. (2019); Neacsu et al. (2022)).

6 http://fcon_1000.projects.nitrc.org/indi/retro/cobre.html

7 https://www.fil.ion.ucl.ac.uk/spm/

8 Recent work has revealed the implication of DMN subsystems in the execution of certain external activities as well (see Mancuso et al. (2022) and the references therein).

9 https://identifiers.org/neurovault.image:568084

10 http://trendscenter.org/software/gift/

11 Note that vanilla spectral DCM is typically used for causal networks with less than 15 regions. For larger networks, further linearized variants of DCM can be used (Frässle et al. (2021); Friston et al. (2021))

12 Ventro-median prefrontal cortex (VMPFC)

13 Antero-median prefrontal cortex (AMPFC)

14 Dorsal prefrontal cortex (DPFC)

15 Temporal pole (TP)

16 Middle temporal gyrus (MTG)

17 Cerebellar hemisphere (CbH)

18 Cerebellar tonsil (CbT)

19 Since cross-spectra are the Fourier counterparts of cross-correlations, spectral DCM is essentially a causal model of how fMRI functional connectivity is generated.

20 Hence the units of *Hertz* for the effective connections encoded in matrix *A*.

21 Blood oxygenation level dependent (BOLD)

22 Self-connections control the region’s excitatory-inhibitory balance, or equivalently its gain or sensitivity to inputs.

23 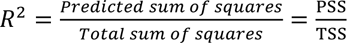, where PSS denotes the sum of squares of cross-spectra predicted/modeled by spectral DCM, and TSS denotes the total sum of squares of cross-spectra estimated from empirical data. The summations (of cross-spectra squared) are performed over frequency bins and pairs of regions for which the cross-spectra are computed. In SPM12, *R*^2^ for any fitted DCM can be calculated using spm_dcm_fmri_check.m.

24 Bayes factor (BF) is the ratio of model evidences (aka marginal likelihoods, ML) of two competing models explaining the same data *y*. That is, BF ≜ P(y|M1)/P(y|M2) = ML1/ML2. In the log space, log BF = log (ML1) – log (ML2) ≍ F1 – F2 = ΔF. So, when ΔF = 3 nats, model 1 is deemed *e*^3^ ≈ 20 times more plausible than model 2, reminiscent of the conventional 0.05 threshold in classical statistics.

25 The complexity term is the KL-divergence between the approximate posteriors and the priors. For the derivations of the group-level free energy of PEB, please refer to (Friston et al. (2015); Friston et al. (2016))

26 In this sense, BMR for DCM is similar to *counterfactual analysis* for SEM Glymour et al. (2016); Pearl (2010).

27 Controllability quantifies the capability of a node/module to drive the dynamical system towards a desired state, using external input. Higher controllability reflects lower average control energy needed to drive the network from that node or set of nodes (Cai et al. (2021); Gu et al. (2022); Liu and Barabási (2016); Pasqualetti et al. (2014)).

28 Causal outflow has been defined as the absolute weighted out-degree of a node on a causal graph (Cai et al. (2021)).

29 Dynamical flexibility refers to the brain’s propensity to *transition* between multiple functional states (Gu et al. (2022)).

## Notes

https://coins.trendscenter.org/

https://identifiers.org/neurovault.image:568084

### Competing Interest Statement

The authors have declared no competing interest.

